# HyperCom: a hypergraph-based method to infer cell-resolved cell-cell communication

**DOI:** 10.64898/2026.09.30.755742

**Authors:** Justin K. Moy, Pawel F. Przytycki

## Abstract

Cell-cell communication underpins intercellular responses allowing cells to dynamically change states across conditions; however current methods typically aggregate signals by cell type annotations which may introduce biases. We developed HyperCom, a hypergraph-based method to infer cell-resolved cell-cell communication from single cell transcriptomics data. HyperCom constructs a hypergraph from known ligand-receptor interactions and the expression of those ligands and receptors in individual cells and performs graph diffusion on the hypergraph to score cells on a continuous spectrum from sender to receiver for each LR pair. HyperCom then generates a cell-type agnostic ranking of LR pairs by communication either over the whole dataset or in a condition-aware manner. Validation of HyperCom on physical cell-to-cell contact, spatial, and multicondition datasets highlight its flexibility to capture subtle per cell communication information. HyperCom is available as a species-agnostic, scalable, and open source R package: https://github.com/pawelab/HyperCom.

## Introduction

Cell-cell communication underpins intercellular responses allowing cells to dynamically change states across conditions. Paracrine signaling, a form of cell-cell communication between nearby cells, is facilitated through ligands which are secreted by cells and bound by receptors on the cell membrane.^1^ Ligands bind to receptor proteins or protein complexes to propagate signals throughout the cell. These interactions can be measured by transcriptomics since many ligands and receptors are translated from RNA.^2^

Single cell transcriptomics (e.g., scRNA-seq) can measure the expression of genes that code for ligands and receptors in individual cells. Several computational tools exist to interrogate cell-cell communication in scRNA-seq. Broadly speaking, these tools rely on a precurated database of ligand receptor (LR) pairs and measure expression of these pairs across cells.^3^ Most tools, including CellChat, CellPhoneDb, and SigXTalk, aggregate gene expression together by cell type.^4–6^ CellChat works by finding differentially expressed genes by cell type and then calculating the mass action of the LR pair for each pair of cell types. CellPhoneDb identifies LR pairs where all components are expressed in at least 10% of a cell type and then finds significant LR pairs by comparing cell type mean expression against a null distribution of permuted cell type labels. SigXTalk uses hypergraph learning to identify communication modules and cross talk between different cell types based on CellChat*’*s top choices. Despite the strengths of these methods, cell type clustering can introduce biases like omitting rare cell populations or discretizing continuous cell states. Alternative non-cell type-based methods like Scriabin can interrogate single pairs of cells, but are computationally infeasible to run on more than a few hundred cells.^7^

To address these issues, we developed HyperCom, a hypergraph-based method to infer cell-resolved cell-cell communication from single-cell transcriptomics data. HyperCom constructs a hypergraph by combining precurated ligand-receptor interactions with the expression of those ligands and receptors in individual cells. Graph diffusion on this hypergraph calculates a global influence matrix which provides the basis of two functionalities: per cell *“*LR Score*”* and dataset wide, per-condition *“*LR Priority*”. “*LR Score*”* assigns a score to cells on a continuous spectrum from sender to receiver for a given LR pair, while *“*LR Priority*”* ranks each LR pair in a dataset by its communication in each condition, timepoint, or other supplied metadata. We benchmarked HyperCom*’*s performance against current tools and demonstrate its capacity to detect up and down regulated LR pairs agnostic of a user-specified control condition, capture signals from experimentally measured physical cell-to-cell contact, identify key biomarkers in lowly expressed spatial transcriptomic data, and distinguish between subtle expression differences in multicondition datasets.

## Results

### Overview of HyperCom

HyperCom interrogates cell-cell communication through network diffusion on a hypergraph, providing two main functionalities: a per cell *“*LR Score*”* and a per dataset *“*LR Priority*”. “*LR Score*”* assigns a continuous score to each cell ranging from sender to receiver for a specific LR pair. *“*LR Priority*”* determines which LR pairs in the dataset have the highest communication either overall or per optional sample metadata such as condition or timepoint (Figure 1). To generate these scores, HyperCom first constructs a heterogeneous hypergraph where nodes represent individual cells, ligand genes, or receptor genes, and hyperedges represent the relationships between these nodes (Figure 1A). Hyperedges, unlike the pairwise edges in a regular graph, encode sets that can contain multiple nodes.^8^ HyperCom uses hyperedges to capture two different relationships: gene expression by cells and LR pair interactions between gene products. Each gene expression hyperedge contains a gene node and all cells that express that gene. LR pair hyperedges contain the genes of a given LR pair to capture the interaction between a ligand and its receptor or receptor complex. Together these nodes and hyperedges form the HyperCom hypergraph.

**Figure 1.**
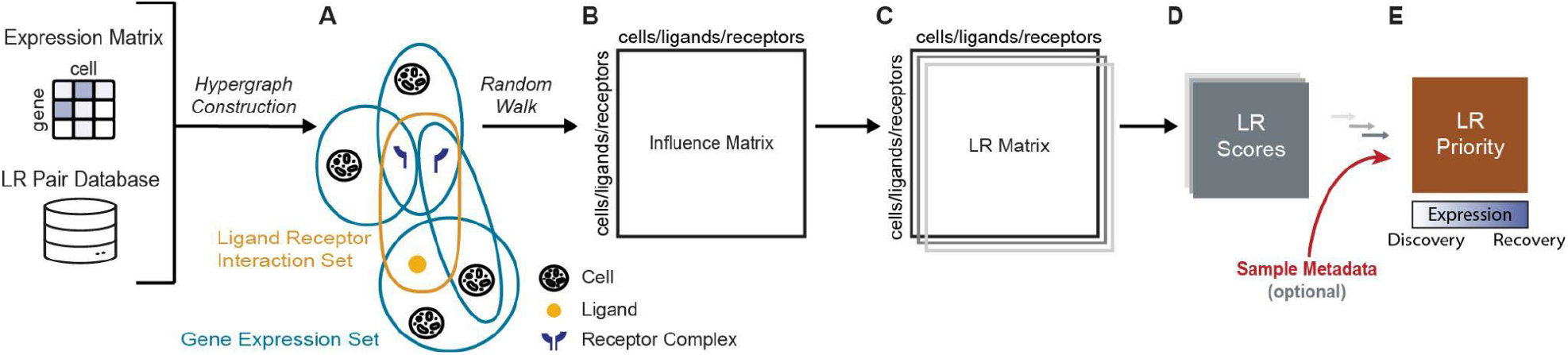
Overview of HyperCom. **A.**HyperCom takes a cell by gene expression matrix and LR pair database as input to construct a hypergraph. **B**. This hypergraph is used to calculate an influence matrix using a random walk. **C**. For each LR pair, an LR matrix is generated from the influence matrix. **D**. LR matrices are used to determine a per cell LR score and **E**. aggregated, optionally using sample metadata, to prioritize LR pairs. LR priority can be tuned to weigh gene expression highly for Recovery Mode or lowly for Discovery Mode.

Although this hypergraph includes observed and known direct relationships between ligands, receptors, and cells, it does not encode the global connectivity of nodes in the network. The network does not inherently capture multistep communications, and due to the sparsity of scRNA-seq, may have missing edges. To address this, HyperCom performs a random walk with restart on the hypergraph to produce an influence matrix (Figure 1B). The influence matrix quantifies the communication from all nodes to all nodes rather than just direct neighbors. A given entry in this influence matrix represents the probability of a walk starting at a row node and then ending at a column node, for example from a ligand to a cell or from a cell to another cell, and column means capture the global average signal that is being received by each cell.

Once constructed, the influence matrix can be used to examine the communication of each cell for a specific LR pair. For each LR pair, HyperCom generates an LR matrix that captures how much cells communicate with each other over that interaction by multiplying the ligand column by the receptor row, then averaging over all parts of the receptor complex (Figure 1C). Taking the row means of this matrix generates a sender score for each cell which represents a cell*’*s affinity for sending signals using the given LR pair and taking the column means of this matrix generates a receiver score for each cell which represents the affinity of a cell to receive signals using the given LR pair. To generate a final *“*LR score*”*, HyperCom takes the difference of normalized sender and receiver scores to produce a single score for each cell (Figure 1D). This HyperCom score ranges from -1 to 1 with -1 representing better senders and 1 representing better receivers for the given LR pair. Since cells with HyperCom scores near 0 can either have low communication or have similarly high sender and receiver scores, HyperCom outputs HyperCom scores, sender scores, and receiver scores per cell. These per cell scores can be used to individually characterize cells, or be aggregated to examine cell-cell communication across groupings of interest like cell types.

Finally, HyperCom can determine which LR pairs, either across a whole dataset or, optionally, in a condition-aware manner, are significant to cell signalling (Figure 1E). This *“*LR Priority*”* score ranks each LR pair using the weighted geometric mean of five terms to account for the expectation that only a small subset of expressed LR pairs will exhibit high communication or condition specificity. The first three terms, which are calculated on the subset of cells expressing a ligand or receptor, are: 1) the mean of the maximum between sender and receiver score of cells to prioritize highly communicative pairs, 2) the gini impurity between the number of sender and receiver cells to enforce ligand and receptor balance, and 3) the gini impurity between the sample conditions to prioritize condition specific interactions. The final two terms, which are calculated using all cells, are: 4) the minimum of the fractions of cells adjacent in the hypergraph to each gene in an LR interaction to ensure both the ligand and receptor are present and 5) the fraction of cells that are adjacent in the hypergraph to any gene in the LR pair to prioritize only LR pairs present in the data. Together these terms ensure that group specific LR pairs with high scoring sender and receiver cells will have higher priority. In order to capture both LR pairs that are highly expressed and LR pairs that are only expressed in a small subset of cells, term five, the fraction of cells adjacent to genes in the LR pairs, can be weighted by a parameter set between 0 for *“*Discovery Mode*”* and 1 for *“*Recovery Mode*”. “*Discovery Mode*”* prioritizes highly communicative but lowly expressed LR pairs like interactions between rare cell populations which are generally missed by expression-based methods. Conversely, *“*Recovery Mode*”* prioritizes highly expressed LR pairs.

### HyperCom simultaneously detects both up and down regulated LR pairs

To assess HyperCom*’*s baseline capacity to identify LR pairs, we tested its ability to recapitulate high confidence interactions from scRNA-seq data predicted by CellChat, the most widely used tool for inferring cell-cell communication per cell type. We ran HyperCom LR Priority in recovery mode on the human skin lesion scRNA-seq dataset containing lesional and nonlesional samples used by Jin et al. to validate CellChat.^9^ Since HyperCom simultaneously prioritizes both up and downregulated LR pairs, we ran CellChat twice to identify each of those types of pairs. To capture how well HyperCom could recapitulate interactions inferred by CellChat, we treated this combined CellChat list as the ground truth on which HyperCom achieved 100% precision at 10% recall and an overall AUPRC of 0.689 (Supplemental Figure 1A) that drops off as expected going from recovery mode to discovery mode (Supplemental Figure 1B).The highest ranking LR pair not detected by CellChat was *TNF*-*TNFRSF1A*. In this case, the ligand was expressed in low quantities by cells classified in five different cell types (Supplemental Figure 2A), while the receptor was primarily expressed in just one (Supplemental Figure 2B). The amount of expression per cell type was not strong enough to be captured by CellChat. However, because HyperCom avoids discrete cell type annotation, it captured subsets of each cell population that when taken together lead to both strong sender and receiver groups (Supplemental Figure 2C). *TNFRSF1A* is an important receptor with regards to inflammation with mutations in this gene shown to cause TNF Receptor-Associated Periodic Fever Syndrome characterized by sudden severe inflammation.^10^ Overall, without any clustering of the cells or annotation of cell types, HyperCom was able to reproduce the inferences made by CellChat and identify communication common across cell types.

HyperCom was able to efficiently infer these interactions. To assess HyperCom*’*s performance we compared time and memory usage against CellChat, which scores cell types as a whole, and Scriabin, which scores individual cells (Supplemental Figure 3).^4,7^ While Scriabin could not be run on more than 1,000 cells, making it unsuitable for typical analysis of single-cell data, HyperCom can be run on 10,000 cells in about 2 hours using 8 cores and 140 GB of memory. Since CellChat aggregates cells by cell type, runtime and memory usage remained constant with increasing cell counts.

### HyperCom recapitulates measurements of physical contact between individual cells

To validate HyperCom*’*s scoring of individual cells*’* communication, we tested its ability to capture physical cell contact as measured by the uLIPSTIC platform.^11^ uLIPSTIC is a sequencing technology that measures cell-cell contact through biotinylation by an engineered donor cell population. Because the biotinylation reaction only occurs if two cells physically touch, the biotin score verifies physical contact at a single cell level. In this protocol, biotin scoring is followed by scRNA-seq which allows for gene expression profiling and cell-contact to be measured in the same cells (Figure 2A).

**Figure 2.**
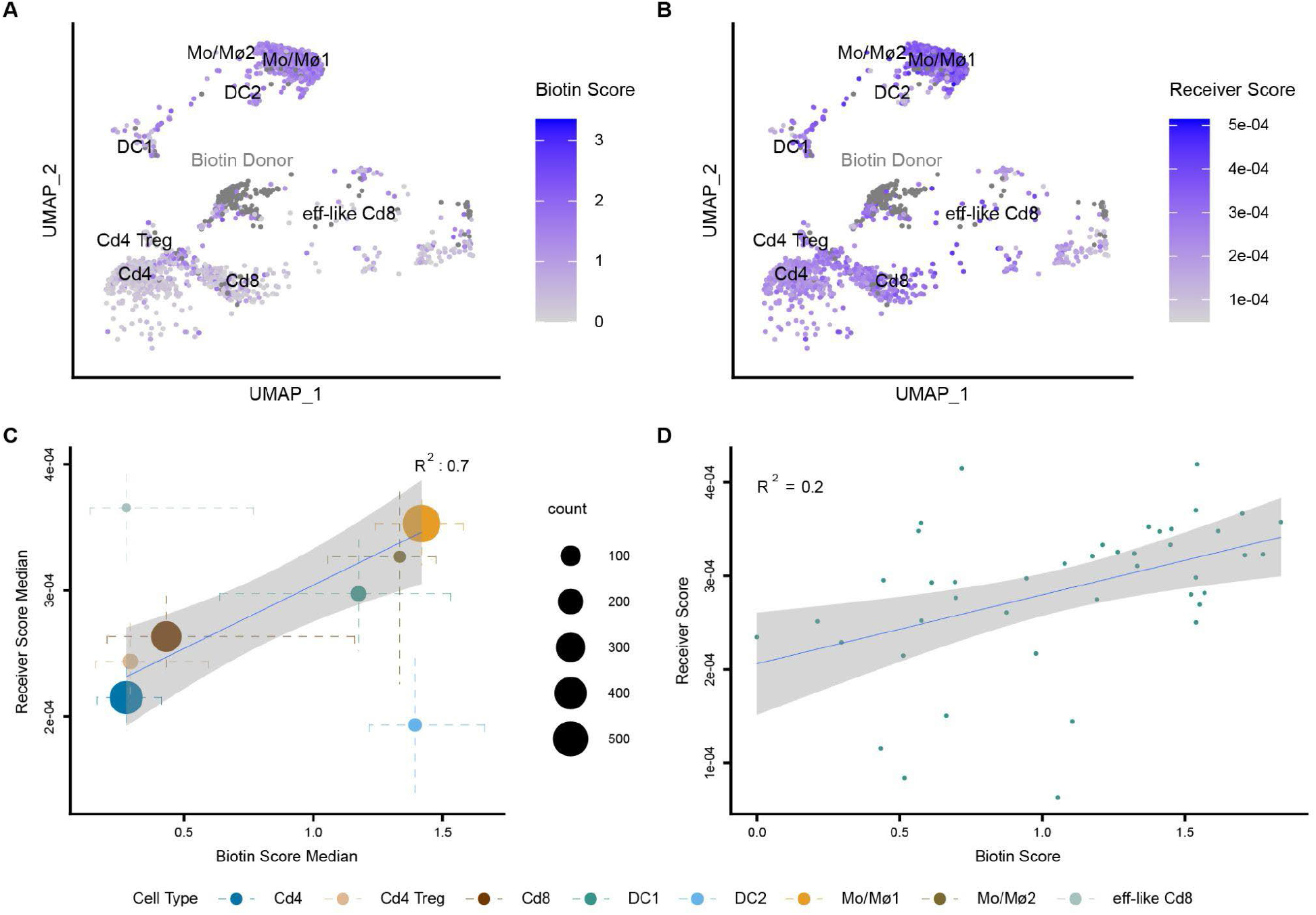
Validation of HyperCom with uLIPSTIC. **A**.UMAP of cells colored by biotin score with donor cells marked with gray. **B**. UMAP of cells colored by HyperCom receiver score with donor cells marked in gray. **C**. Regression of biotin score against receiver score for each cell type. Each point is placed at the median biotin score and median receiver score for cells in that cell type with size scaled by the number of cells. The bars for each point show the interquartile range for each cell type for each score. The blue line shows linear regression fit weighted by the number of cells for each cell type with the gray area showing the 95% confidence band. **D**. Regression of biotin score against receiver score for DC1 cells. The blue line shows linear regression fit with gray area showing the 95% confidence band.

We ran HyperCom LR Priority in recovery mode on uLIPSTIC scRNA-seq data of immune cells in the mediastinal lymph node 36 hours post lymphocytic choriomeningitis virus (LCMV) infection to identify key LR pairs specific to immune response. HyperCom*’*s LR pair ranking overlapped highly with CellChat*’*s inferred interactions with a 100% precision at 10% recall and an overall AUPRC of 0.784 (Supplemental Figure 1C). HyperCom*’*s top ranked LR pairs included many components of the MHC complex including all six pairings of the ligands *H2-K1, H2-D1*, and *H2-Q7* with the receptors *Cd8a* and *Cd8b1*, all of which were also identified by CellChat.^12^ These MHC components showed communication patterns typified by the *H2-K1-Cd8a* interaction where sender cells were not limited to a single cell type (Supplemental Figure 4A), but only Cd8 cells were receivers (Supplemental Figure 4B). Additionally, among the top interactions, HyperCom significantly ranked the *Cd86-Cd28* interaction, which is known to modulate T cell activity, with higher sender scores for DC1 cells and monocyte/macrophages (Supplemental Figure 4C) and high receiver scores for Cd8 and Cd4 cells (Supplemental Figures 4D).^13^

Given that HyperCom can recapitulate the per cell type results of CellChat on this dataset, we next examined HyperCom*’*s ability to capture known physical contact at single cell resolution. Using the previously annotated labels of donor and acceptor cells based on biotinylation, we calculated the global average signal received using only donor cells as senders (Figure 2B). A regression of cell type median biotin score against median receiver score weighted by the number of cells in each cell type had a strong positive correlation (Figure 2C, R^2^ = 0.70, *p-*value = 0.0027) as did per-cell scores (Supplemental Figure 4E, R^2^ = 0.24, *p*-value < 2e-16).

Of particular interest are DC1 cells which are critical for initial immune response and are expected to ramp up communication as infection progresses.^11^ We tested if HyperCom was able to detect this gradual change by performing a per cell regression between HyperCom receiver score and biotin score for just DC1 cells which yielded a significant positive correlation (Figure 2D, R^2^ = 0.2, *p*-value = 0.0029). This ramp up of antigen presentation is obscured by methods that detect cell-cell communication via cell type aggregation and contrasts with the ubiquitous communication in macrophages and naive Cd8 cells and ubiquitous lack of communication in Cd4 cells (Supplemental Figure 4F).

### HyperCom captures spatial communication patterns

Spatial transcriptomics captures both the location and quantity of gene expression in a tissue. Location measurements of spatial transcriptomics can proxy cell-cell communication since proximal cells tend to communicate more with each other than distal cells.^14,15^ Therefore to further assess HyperCom*’*s ability to identify interactions, we ran HyperCom on a high resolution section of ovarian cancer tissue with a large diversity of cell types processed with the Xenium imaging-based spatial transcriptomic platform, capturing over 5000 genes (Figure 3A).^16^ As expected, the influence matrix values between pairs of cells had a weak but highly significant negative correlation with Euclidean distance between cells (Pearson*’*s *r* = -0.021, *p*-value < 2e-16) indicating that HyperCom successfully captures the proclivity of proximal cells to communicate more.

**Figure 3.**
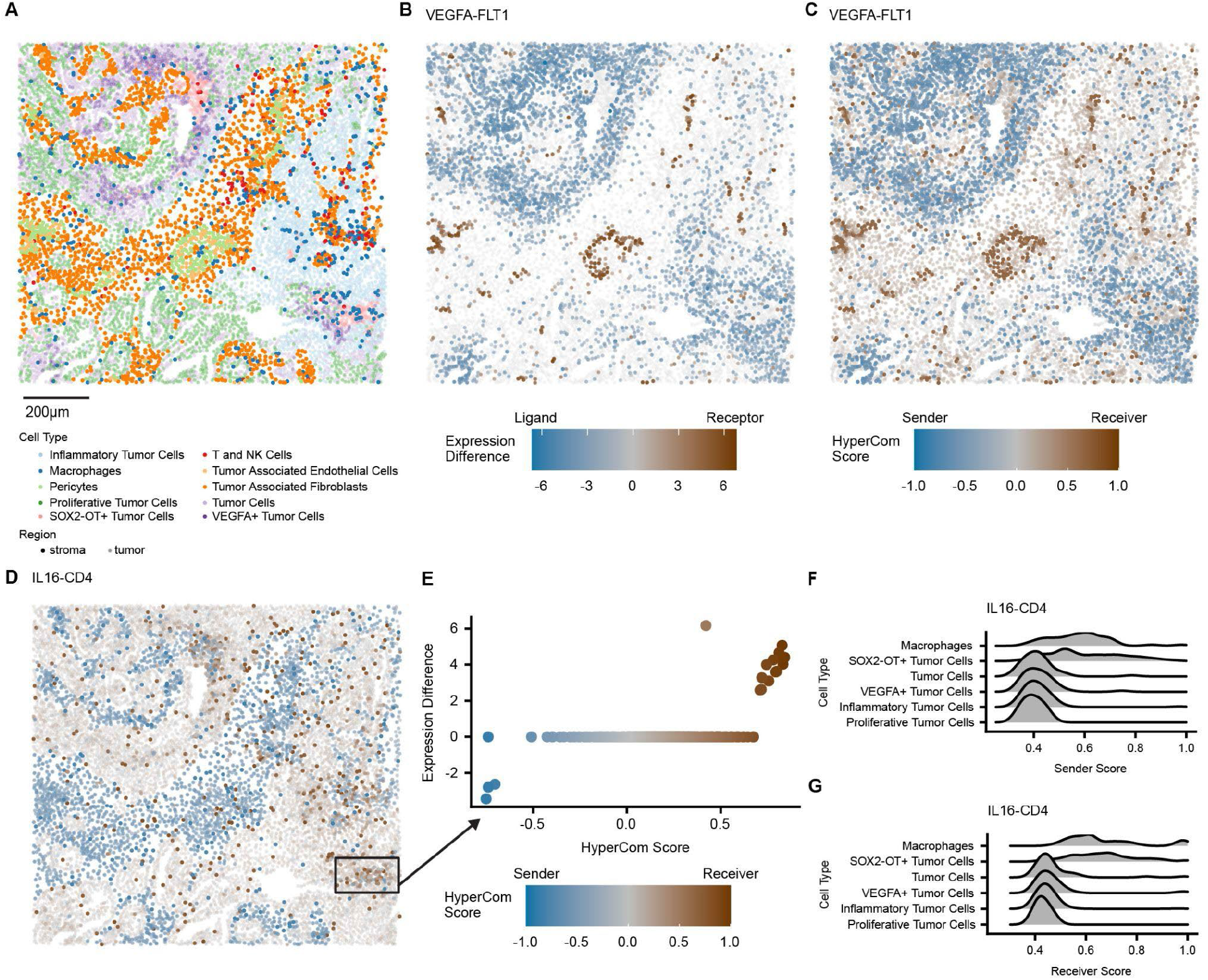
HyperCom Detects Spatial Communication. **A**.Spatial locations of cells colored by region and annotated cell type. **B**. Spatial locations of cells colored by difference in VEGFA (ligand) and FLT1 (receptor) expression. Blue represents higher ligand expression and brown represents higher receptor expression. The alpha is set to the maximum between the ligand and receptor expression. Spatial locations of cells colored by HyperCom score for **C**. VEGFA-FLT1 and **D**. IL16-CD4. Blue indicates higher sender scores and brown indicates higher receiver scores. The alpha is set to the maximum between the sender and receiver score. **E**. Scatter plot of HyperCom score versus expression difference for cells in the highlighted region in D. Joy plots split by cell type for cell types with at least 5 cells in the highlighted region in D sorted by mean score for **F**. IL16-CD4 sender scores and **G**. IL16-CD4 receiver scores.

To test HyperCom*’*s overall ability to capture communicating LR pairs from spatial data, we first ran it in recovery mode and achieved 98.3% precision at 10% recall and an overall AUPRC of 0.530 for recovering pairs found by CellChat (Supplemental Figure 1D). Among the top significant pairs was the *VEGFA-FLT1* interaction, in which the ligand and receptor were both highly expressed (Figure 3B) and HyperCom scores mirrored the expression pattern (Figure 3C, Pearson*’*s *r* = 0.940, *p*-value < 2e-16). This well-established interaction facilitates angiogenesis whereby cancer cells express *VEGF* and pericytes, which form blood vessels, express *FLT1*, also known as *VEGFR1*.^*17,18*^ Concordantly in this dataset, *VEGFA+* tumor cells had a high mean sender score (Supplemental Figure 5A), and pericytes had a high mean receiver score score (Supplemental Figure 5B).

We next ran HyperCom in discovery mode, which identifies lowly expressed interactions. Among the top ranked significant interactions not identified by CellChat or HyperCom in recovery mode was *IL16-CD4*. In ovarian cancer, *IL16*, a cytokine that promotes angiogenesis and immune recruitment through its binding to *CD4*, has been proposed as a potential biomarker for poor ovarian cancer prognosis.^19,20^ However, in this sample, of 9918 cells, only 216 were detected to express *IL16* and 301 *CD4*. Despite this low expression, HyperCom finds spatially co-localized senders and receivers (Figure 3D). A close inspection of one such region shows a dynamic range of HyperCom scores even for cells with almost no detected gene expression (Figure 3E). Breaking down HyperCom scores by cell type shows communication between SOX2-OT+ tumor cells and macrophages (Figure 3F and G).

### HyperCom distinguishes communication between multiple conditions

To showcase HyperCom*’*s functionality on multicondition data, we explored cell-cell communication in a two-condition dataset of the transverse aortic constriction (TAC) model of murine chronic heart failure and a three condition dataset of immune cells from seven days post TAC surgery with *Brd4* knockout.^21^ The first dataset includes cells collected from the hearts of healthy mice (Sham) and mice sixty days post TAC surgery. We ran HyperCom on this data in recovery mode to identify condition-specific cell-cell communication and benchmarked it against CellChat. Since HyperCom detects communication across conditions simultaneously, in order to generate a comparable set of interactions, CellChat had to be run twice, first to detect LR pairs up in TAC and then interactions up in Sham. HyperCom achieved 81.5% precision at 10% recall and an overall AUPRC of 0.424 on the full set of LR pairs identified by CellChat (Supplemental Figure 1E). Highlighting HyperCom*’*s ability to detect interactions across conditions, HyperCom*’*s top LR pair by priority score was *Thbs1*-*Sdc4*, which is enriched in TAC (Figure 4A) while the second ranked LR pair was *Pgf*-*Flt1*, which is enriched in Sham (Figure 4B). Although HyperCom scoring is cell type independent, breaking down the scores by annotated cell type revealed that in the first interaction, signal was primarily sent by fibroblasts and received by macrophages (Figure 4C and D). This interaction is a known stress response with *Thbs1* implicated as an activator of the fibrotic TGF-β pathway and cardiac hypertrophy.^22,23^ In the second interaction, senders were primarily enriched in fibroblasts while receivers were enriched in both endothelial and lymphatic endothelial cells as shown by mean sender and receiver score (Figure 4E and F). *Pgf*-*Flt1* has been shown to play a role in angiogenesis and healing after cardiac stress.^18,24,25^ Both of these interactions were detected by a single run of HyperCom without the need to cluster the data into cell types.

**Figure 4.**
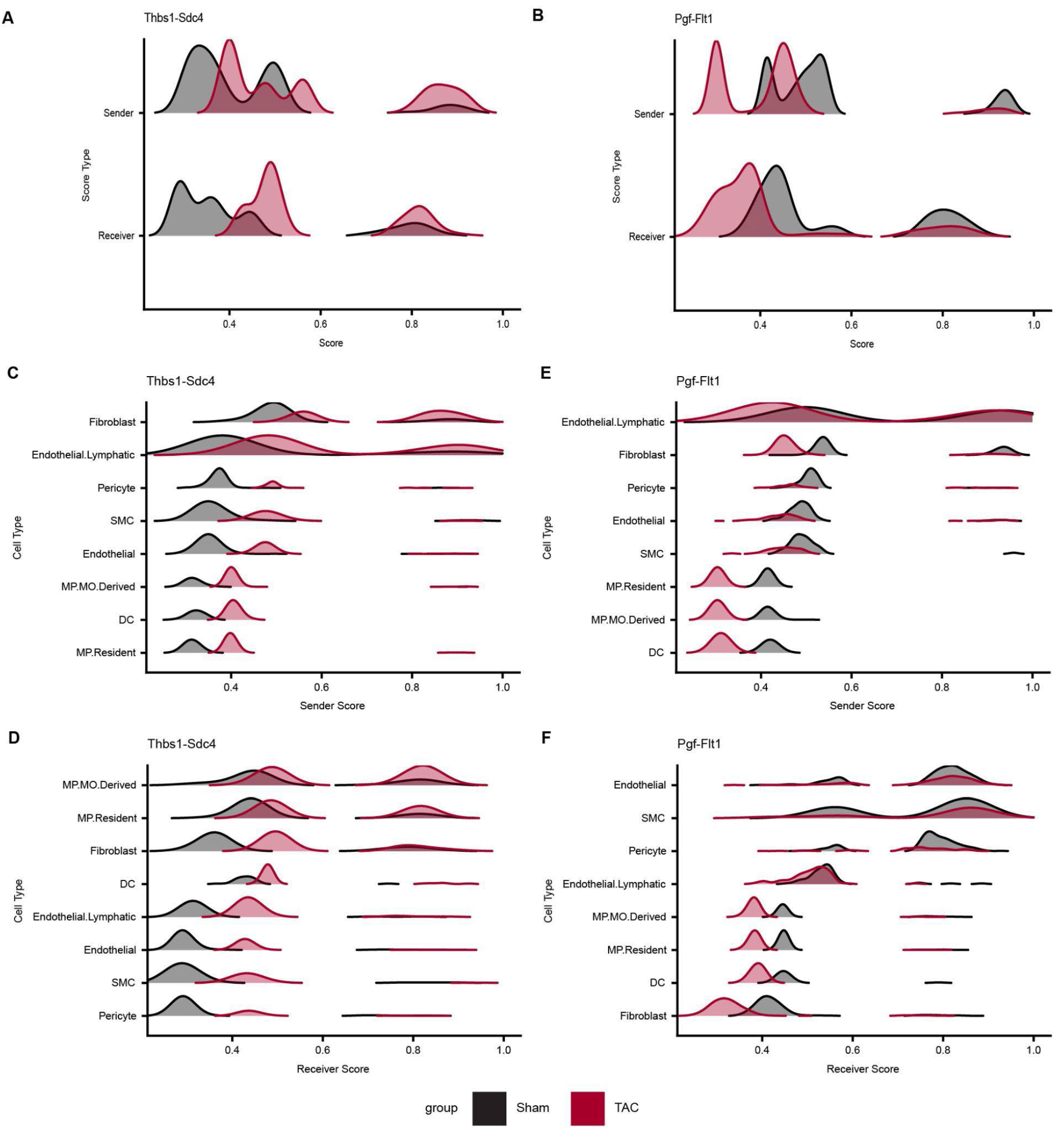
Significant LR Pairs in murine model of chronic heart failure. Joy plots colored by condition for **A**. Thbs1-Sdc4 and **B**. Pgf-Flt1 and grouped by cell type for cell types with at least 50 cells for **C**. Thbs1-Sdc4 sender scores, **D**. Thbs1-Sdc4 receiver scores, **E**. Pgf-Flt1 sender scores, and **F**. Pgf-Flt1 receiver scores.

Finally, we tested HyperCom*’*s ability to detect shifts in communication across three conditions using scRNA-seq immune cell data from seven days post sham surgery, TAC surgery, and TAC surgery with *Brd4* knockout (BRD4KO). *Brd4* is a key regulator of cardiac stress response and inhibition of *Brd4* has been shown to improve heart function in TAC mice.^26^ As such, expression in BRD4KO mice for some genes is expected to return toward baseline Sham levels. While inference on this dataset can be performed with a single run of HyperCom, CellChat would need to be run combinatorially for each pair of conditions, limiting its statistical power. Using recovery mode, *Spp1-Cd44* was the top significant interaction overall and showed an increase in TAC and decrease in *Brd4* knockout (Figure 5A). Similarly, discovery mode prioritized the canonical neutrophil recruitment interaction *Cxcl1-Cxcr2* as significant despite their low expression levels (Figure 5B).^27,28^ These interactions were primarily detected between macrophages and neutrophils (Figure 5C-F), in agreement with their observed role and communication pattern regarding neutrophil recruitment.^29,30^

**Figure 5.**
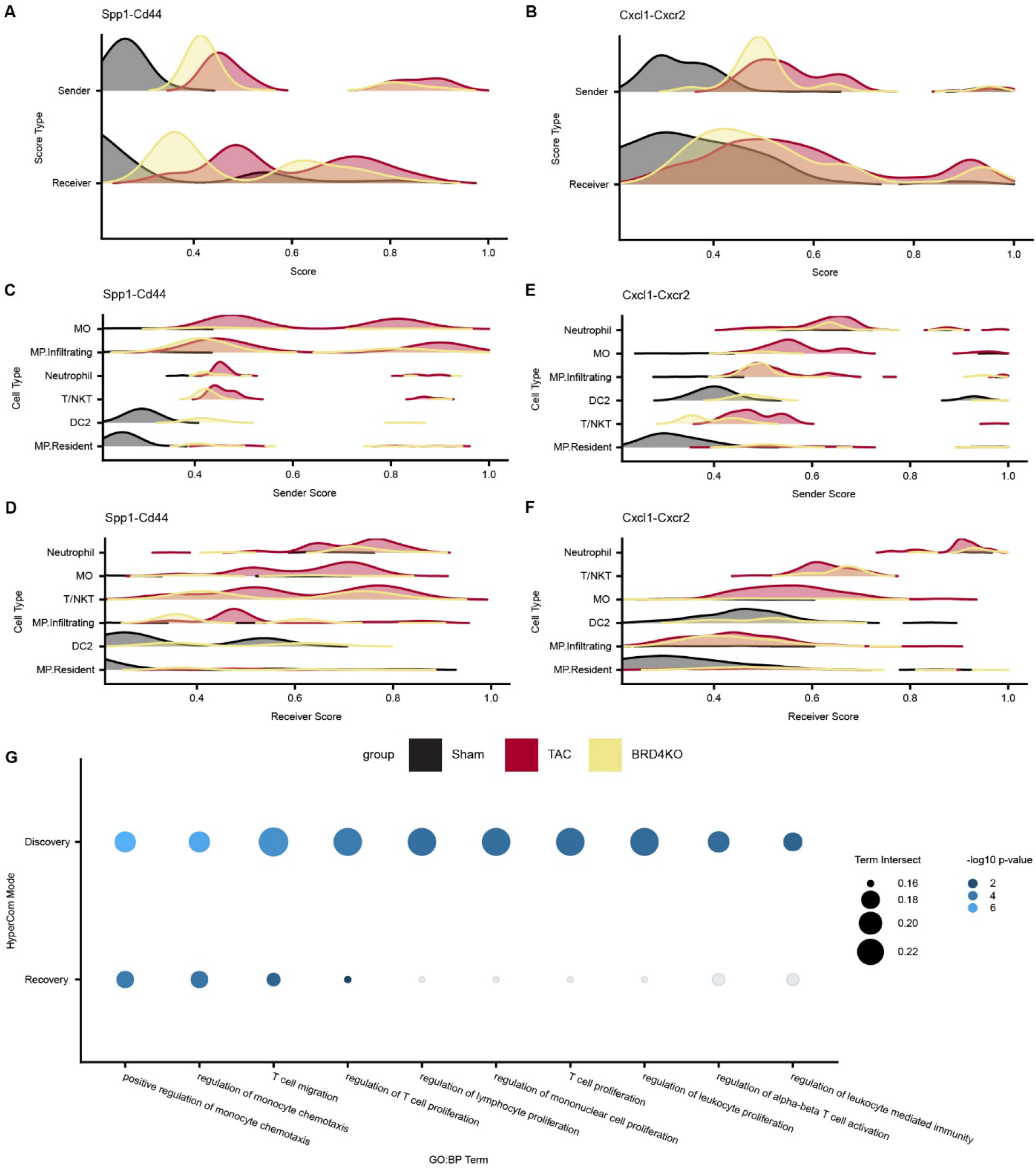
HyperCom detects subtle communication shifts. Joy plots colored by condition for **A**. Spp1-Cd44 and **B**. Cxcl1-Cxcr2 and grouped by cell type for cell types with at least 50 cells for **C**. Spp1-Cd44 sender scores, **D**. Spp1-Cd44 receiver scores, **E**. Cxcl1-Cxcr2 sender scores, and **F**. Cxcl1-Cxcr2 receiver scores. **G**. From left to right, top 10 GO:BP terms ordered by largest decrease in significance from discovery to recovery mode from among significant LR pairs for genes in LR pairs with absolute mean HyperCom scores increasing from Sham to TAC and decreasing from TAC to BRD4KO, scaled by the size of the intersection with the GO:BP term and colored by -log10 p-value for discovery and recovery mode.

To characterize HyperCom*’*s overall ability to capture subtle changes to communication patterns, we performed gene set enrichment on ligand and receptor genes belonging to significant LR pairs grouped by communication patterns across conditions. Out of the 43 total significant LR pairs across recovery and discovery mode, 24 pairs showed an increase in absolute mean HyperCom score from Sham to TAC and decrease from TAC to BRD4KO and 14 showed an increase from Sham to TAC but no decrease in BRD4KO (all other combinations had fewer than 10 LR pairs). The top Gene Ontology Biological Processes (GO:BP)^31,32^ terms largely overlapped for all modes, with the increase/decrease gene set highlighting mononuclear cell movement and monocyte chemotaxis, while the increase/no decrease gene set contained SMAD protein signal transduction, a part of the TGF-B stress response pathway, a known stress response to TAC surgery (Supplementary Figure 6A-C).^33,34^ In order to examine terms derived from rare and lowly expressed interactions, we sorted significant terms by largest decrease in significance from discovery to recovery mode. This approach highlighted terms involved in leukocyte and T cell immunity in the increase/decrease gene set (Figure 5G), while the increase/no decrease gene set featured terms related to natural killer cells (Supplemental Figure 6D). HyperCom was able to identify these subtle condition-specific shifts due to its ability to simultaneously interrogate multiple conditions even for lowly expressed LR pairs.

## Discussion

HyperCom interrogates cell-cell communication at a truly single cell resolution unbiased by cell type annotation. The hypergraph construction and subsequent random walk support the inference and scoring of LR pairs that may be obscured by low or no detectable LR expression. LR scoring provides the granularity to examine the communication of individual cells across specific LR pairs through either sender, receiver, or a single HyperCom score per cell. Additionally, these scores can still be aggregated to examine biological groupings of interest like cell types. HyperCom*’*s LR priority then highlights LR pairs of interest, and can simultaneously prioritize LRs across multiple conditions in a single run. Additionally recovery and discovery mode capture both highly and lowly expressed interactions that may not be captured from expression alone.

HyperCom*’*s hypergraph-based design makes it a flexible method with the potential to capture heterogeneous data types. In this work, we chose to binarize expression as well as treat ligand and receptor complex genes with equal importance during hypergraph construction. This reduced computation and maintained interpretability without making assumptions on specific biological thresholds for expression. Through this approach, HyperCom captured weak interactions, recapitulated gene expression levels, and identified spatial patterning. However, edges in the graph have the potential to be weighed by factors such as expression level and spatial distance between cells, and new hyperedges could be added to capture regulatory relationships and other modalities such as chromatin accessibility.

While most methods for detecting cell-cell communication are only able to do so at a cell type level, HyperCom scores individual cells and is able to do so much more efficiently than existing cell-resolved methods. However, while HyperCom can score tens of thousands of cells, modern cell atlases reach into the hundreds of thousands or even millions of cells that span multiple different individuals, collection sites, and conditions.^35^ The growth of atlases is outpacing increases in computational speed. HyperCom has the potential to be applied at these scales by operating on a sample of representative cells or aggregated metacells, both of which HyperCom can readily be adapted to.^36,37^

Like all current cell-cell communication methods, HyperCom fundamentally depends on a supplied database of LR pairs. For species like mice and humans there exists a wide array of databases and annotations, but other less studied organisms*’* precurated interactions may be sparser. HyperCom itself can be run for any species, but if an interaction is not present, it will not be taken into account during analysis.

HyperCom is species agnostic, scalable, and easy to install and run through an open source R package. HyperCom*’*s design provides the flexibility to run key steps with intermediate outputs that gives researchers options when rerunning analyses rather than starting from scratch. Additionally the LR priority function lets users choose how much they weigh expression allowing for not only recapitulating expression, but also discovering novel experimental avenues. Taken together, HyperCom is a powerful tool to understand cell-cell communication at a truly single cell resolved level.

## Methods

### HyperCom

#### Hypergraph Construction

HyperCom takes as input a gene by cell expression matrix and a list of ligand-receptor interactions from a ligand-receptor database such as CellChatDB.^4^ First the matrix is filtered to only retain cells expressing ligand and receptor genes from the database with cases where the ligand and receptor are the same gene omitted. A hypergraph is then constructed with cells, ligands, and receptors as nodes and with hyperedges representing two types of interactions: gene expression and ligand-receptor interactions. For gene expression hyperedges, for each gene, cells that have any measured expression of that gene are added to the expression hyperedge of that gene. Interaction hyperedges are built as sets of receptors and ligands that interact either pairwise or as a complex. The full hypergraph is the concatenation of these two edgelists.

#### Random Walk

In a random walk on a hypergraph, a walker can transition from any node in a hyperedge set to any other node in the set. To generate the corresponding row-stochastic transition matrix *W* we assign an equal transition probability between all nodes within a set. Then the steady state global influence matrix *F* is calculated as random walk with restart as *F = α(I − (*1 *− α*)*W*)^*−*1^ where *I* is the identity matrix and *α* is the random restart probability. Alpha is set to 0.5 by default as other network diffusion methods have found that value captures graph structure while still traversing the graph effectively.^38,39^ The influence matrix *F* is also row-stochastic with each row representing the probability of a walk starting at that node and ending at each other node.

#### LR Matrix Generation

To calculate the communication of all pairs of cells over a specific LR pair, the influence matrix is subset to the cell-by-cell portion and the outer product of the column of the ligand gene is taken with the row of the receptor gene. Here the column represents every cell going to the ligand and the row represents the receptor going to every cell. The outer product produces a square matrix representing a score for each cell going to every other cell in an LR-pair specific context. This LR-pair matrix is then scaled through element-wise division by the influence matrix. For ligands with a receptor complex, the matrix is calculated for each component, then all ligand-component matrices are averaged element-wise. Finally, entries of this matrix between cells that come from different samples are set to 0.

#### LR Score Calculation

To calculate the LR score for each cell, the LR matrix is used to determine the propensity of a cell toward being a sender or a receiver. Per cell sender scores are calculated as the percentile normalized row means of the LR matrix and receiver scores as the percentile normalized column means of the LR matrix. To create a single score for each cell that captures both the magnitude and direction of cell communication, the HyperCom score is calculated as the percent rank of the difference between sender and receiver score is multiplied by the sign of the difference and the average of the sender and receiver score. This final score ranges from -1 to 1 where -1 represents strong senders and 1 represents strong receivers.

#### LR Priority Calculation

In order to rank the communication of LR pairs either in a dataset as a whole or by optional metadata (timepoint, condition, etc), HyperCom calculates LR Priority as a weighted geometric mean of five terms. The first three terms use only cells expressing the given ligand or receptor:

1. The average, across cells, of the maximum between the sender and receiver score for each cell. This ensures highly communicative pairs are ranked higher.
2. The gini impurity between the number of cells with positive HyperCom scores (senders) and the number of cells with negative HyperCom scores (receivers). This ensures the LR pair is balanced for senders and receivers.
3. The gini impurity between the number of cells in each label of interest in the metadata. This prioritizes LR pairs that vary across labels. The remaining two terms use all cells:
4. The minimum of the fractions of cells adjacent in the hypergraph to each gene in an LR interaction. This ensures both the ligand and receptor (or all genes in the receptor complex) are expressed.
5. The fraction of cells that are adjacent in the hypergraph to any gene in the LR interaction. This ensures that only LR pairs expressed in the data are ranked highly.

By default all five of these terms are weighted equally; however, to identify communication observed in a smaller subset of cells either due to low expression levels or the rarity of the communicating cell population, the fifth term can be weighted between 0 (discovery mode) and 1 (recovery mode) with lower weights putting less emphasis on directly observed expression of the given ligand and receptor genes.

A permutation test is conducted to assess the significance of LR priority scores. Ligand and receptor genes are randomly sampled to form a background set of LR interactions. By default, 1000 of these pairs are created. Priority scores for these LR interactions are calculated and the fraction of these permuted interactions with scores greater than the score of the true interaction is calculated to generate an empirical *p*-value. To control for multiple-hypothesis testing, *p-*values are then adjusted with Benjamini-Hochberg correction and called significant at an FDR threshold of 0.05.

#### Data Processing

The scRNA-seq datasets we used in this study were processed using Seurat (version 5.3.0) as follows:^40^

The human skin skin lesion RDA file was downloaded from the CellChat vignette.^41^ The gene expression count matrix was extracted from the data slot of the RDA object and the metadata was extracted from the meta.data slot of the object. Together the count matrix and metadata were combined into a Seurat object using the CreateSeuratObject function from the Seurat package. The final Seurat object contained eight samples: four for the nonlesion condition (samples 1,2,5, and 7) and four samples for the lesion condition (samples 3, 11, 14, and 15).

The h5ad LCMV scRNA-seq data from uLIPSTIC was sent by the authors upon request.^11^ The *read_h5ad* function from the anndata package (version 0.7.5.6) to convert the object into a format usable by R.^42^ The gene expression count matrix was extracted from the X slot of the anndata object and the metadata was extracted from the obs slot of the object. Together the count matrix and metadata were combined into a Seurat object using the *CreateSeuratObject* function from the Seurat package. For analysis, the Seurat object was subset to just cells from the 36h condition mouse m11.

The 10X*’*s FFPE Human Ovarian Cancer with 5K Human Pan Tissue and Pathways Panel plus 100 Custom Genes was downloaded from 10X.^16^ The data was loaded into Seurat using the *LoadXenium* function from the Seurat package. The x and y coordinates from the fov slot were then added to the meta.data slot. The data was subset to a 1000 by 1000 slice from X coordinate 1300 to 2300 and Y coordinate 3000 to 4000 containing about 10,000 cells. This slice was chosen due to its diversity of cell types as annotated by 10X.

The day 60 chronic murine heart failure RDA file was obtained upon request from the authors.^21^ The Seurat v2 object was converted to a Seurat v5 using the *UpdateSeuratObject* function from the Seurat package. For analysis, the Seurat object was subset to just cells from samples 2 (Sham) and 3 (TAC).

The day 7 acute murine heart failure RDA file was obtained upon request from the authors.^43^ The Seurat v2 object was converted to a Seurat v5 using the *UpdateSeuratObject* function from the Seurat package. Only cells with nFeature_RNA > 250, nFeature_RNA < 2500, and percent.mt < 5 were retained. The final Seurat object contained five samples: one for Sham (sample 1), two for TAC (samples 2 and 3), and two for BRD4KO (samples 6 and 7).

#### HyperCom Analysis

To calculate LR scores and LR priority, HyperCom was run on each dataset with default settings and discovery mode and recovery mode used as specified for each result. For LR priority calculations, HyperCom was run on the LCMV 36 hr and ovarian cancer datasets without any sample metadata. The skin lesion, day 60 heart failure, and day 7 heart failure datasets were run with condition as the labels of interest from the metadata (lesion/nonlesion, Sham/TAC, and Sham/TAC/BRD4KO respectively).

#### CellChat Analysis

CellChat (version 2.1.2) was run on the skin lesion, LCMV infection, Xenium ovarian cancer, and day 60 heart failure data using default parameters. For the skin lesion and 60 day heart failure data CellChat was run on each condition separately (lesion/nonlesion and Sham/TAC respectively) with upregulated LR pairs in each condition identified by following the CellChat multi-condition vignette. In order to compare HyperCom to CellChat for each of those datasets, the final LR significance table from CellChat was aggregated across cell types using the dplyr (version 1.2.1) *summarize* function by LR pair taking the max communication probability for each pair. The precision recall curve was calculated for HyperCom*’*s LR score against called CellChat LR pairs using the PRROC (version 1.4) package in R.^44^

#### Pathway Enrichment Analysis

Pathway enrichment on day 7 murine heart failure was performed using the gprofiler2 (version 0.2.4) R package.^45^ Significant LR pairs were first identified in a range of modes from discovery to recovery (0, 0.25, 0.5, 0.75, and 1) and then classified either as increasing from Sham to TAC and decreasing from TAC to BRDKO or increasing from Sham to TAC with no decrease from TAC to BRDKO based on the mean of the absolute values of HyperCom scores for the LR pair. Gene set enrichment was run on the genes from the significant LR pairs in each of the 10 combinations of LR priority mode and communication pattern classification using gprofiler2 with the *ordered_query* argument of function *gost* set to TRUE and the genes ordered by the corresponding LR priority mode based on the highest ranking LR pair they appear in. All GO:BP terms with more than 5 and fewer than 200 genes were tested and called significant at an adjusted *p*-value < 0.05.^31,32^

### Implementation

The HyperCom package and all analyses were implemented in R version 4.4.0. The HyperCom package and a vignette detailing an example use case are available at https://github.com/pawelab/HyperCom.

## Acknowledgements

We thank Michael Alexanian for sharing their day 7 murine heart failure dataset and Yuri Pritykin for sharing the LCMV infection dataset.

## Author Contributions

J.K.M. and P.F.P conceptualized the method, J.K.M. performed the analysis, J.K.M. and P.F.P. wrote the manuscript, and P.F.P supervised the study. All authors read and approved the final manuscript.

