## Supplemental Figures for "HyperCom: a hypergraph-based method to infer cell-resolved cell-cell communication"

### Supplement

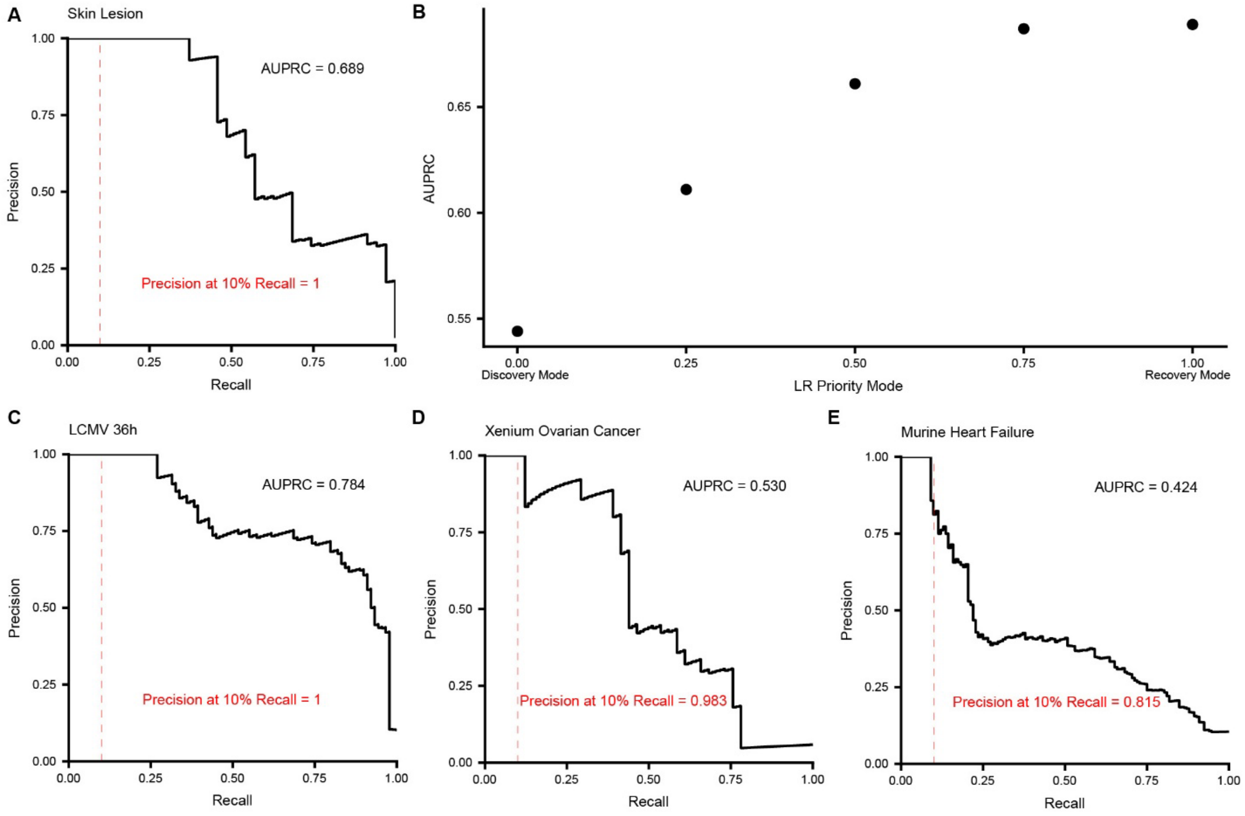


*Supplemental Figure 1.* ***Precision-Recall Curves****.* ***A****. Precision versus recall of HyperCom LR priority in recovery mode for CellChat inferred LR pairs for skin lesion data.* ***B****. AUPRC versus LR priority mode for skin lesion data. Precision versus recall of HyperCom LR priority in recovery mode for CellChat inferred LR pairs for* ***C****. LCMV 36 hours post infection,* ***D****. Xenium ovarian cancer, and* ***E****. 60 day chronic murine heart failure.*

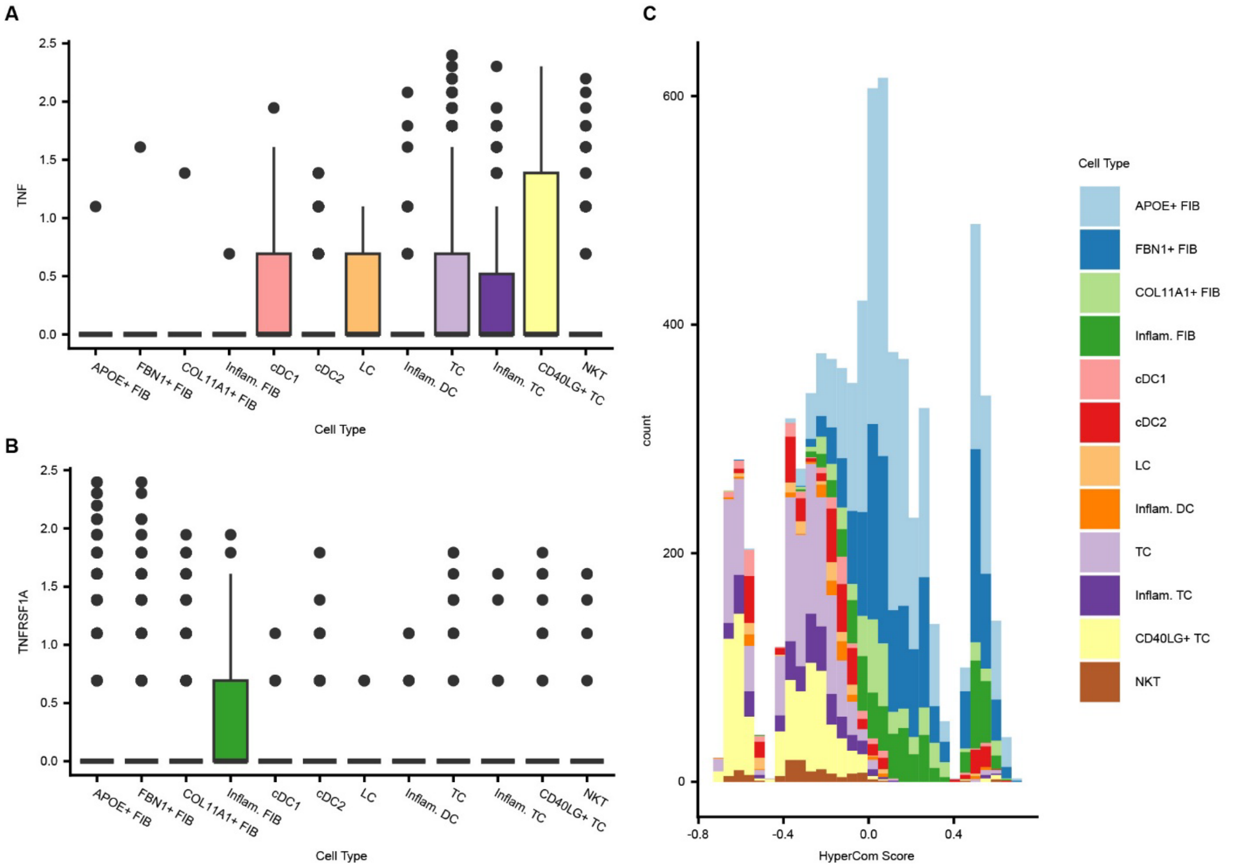


*Supplemental Figure 2.* ***Skin lesion TNF and TNFRSF1A expression and HyperCom scores****. Boxplots of expression per cell by annotated cell type for* ***A****. TNF and* ***B****. TNFRSF1A.* ***C****. Stacked histogram of HyperCom scores colored by cell type.*

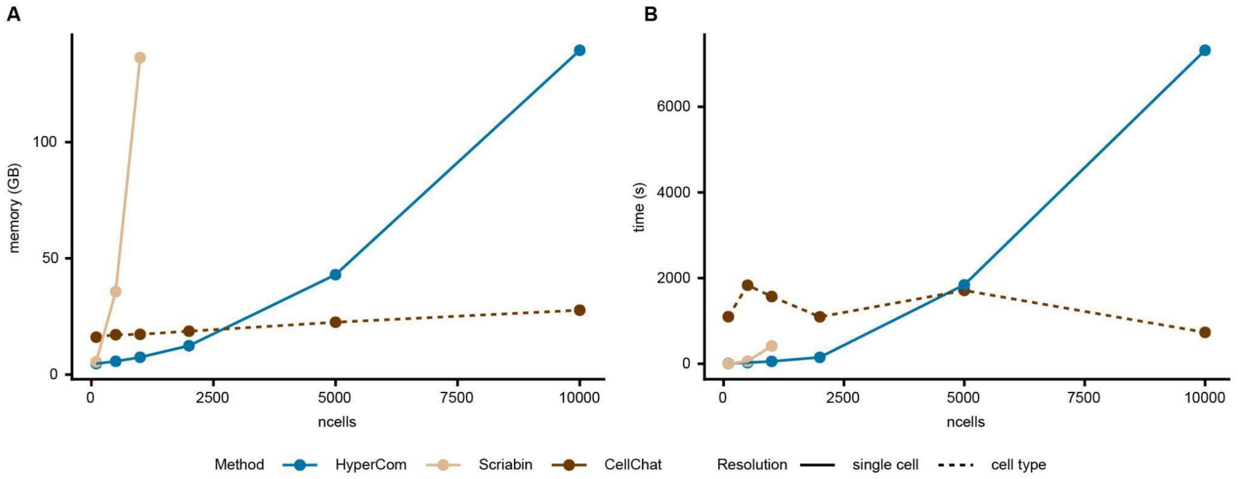


*Supplemental Figure 3.* ***Benchmarking HyperCom performance****. Comparison of* ***A.*** *maximum memory usage (in gigabytes) and* ***B.*** *total run time (in seconds) of HyperCom, Scriabin, another cluster free cell-cell communication method, and CellChat run on random samples of 100, 500, 1000, 2000, 5000, and 10000 cells using 8 Intel Gold cores. Scriabin was run using the cell-cell interaction matrix workflow described in the Scriabin vignettes after optional ALRA denoising.*[7] *Scriabin could only be run for up to 1000 cells due to its high memory usage.*

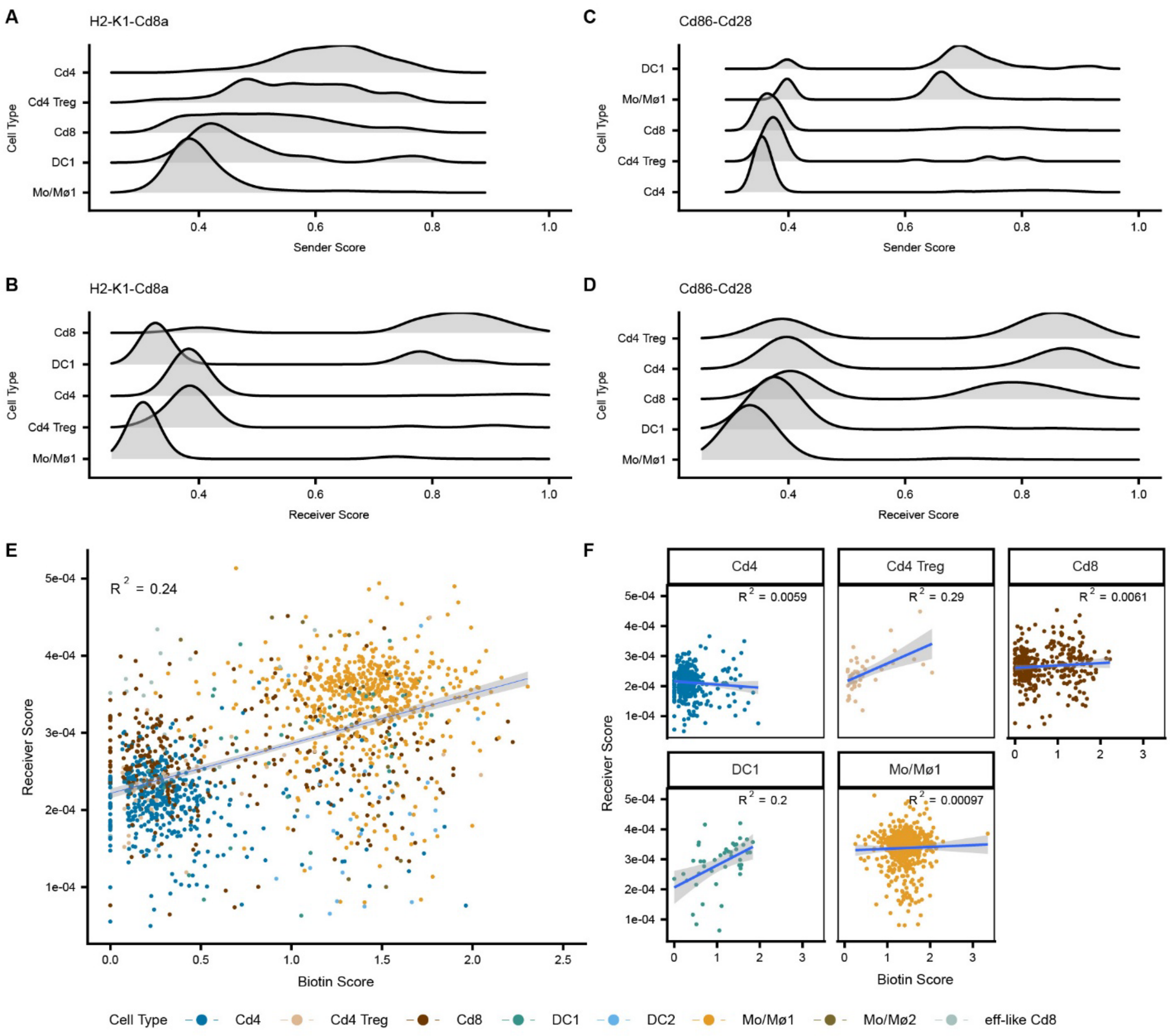


*Supplemental Figure 4.* ***LCMV infection sender and receiver scores****. Joy plots of scores grouped by annotated cell type for cell types with at least 30 cells sorted by mean for* ***A.*** *H2-K1-Cd8a sender scores,* ***B.*** *H2-K1-Cd8a receiver scores,* ***C****. Cd86-Cd28 sender scores, and* ***D****. Cd86-Cd28 receiver scores. Regression of biotin score against receiver score for* ***E.*** *all cells, colored by cell type, and* ***F.*** *grouped by annotated cell types for cell types with at least 30 cells. The blue line shows linear regression fit with gray area showing the 95% confidence band.*

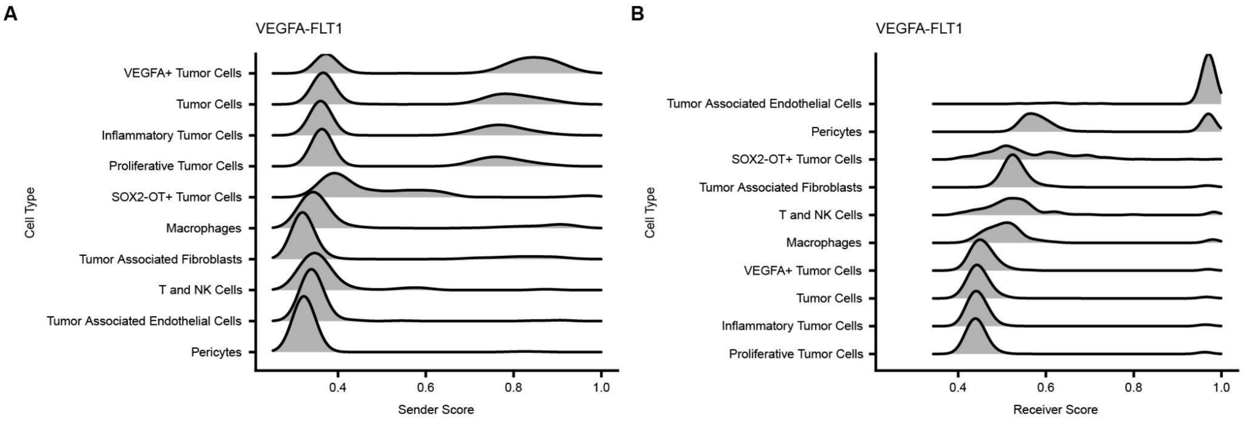


*Supplemental Figure 5.* ***VEGFA-FLT1 sender and receiver scores by cell type in ovarian cancer****. Joy plots of scores grouped by annotated cell type for cell types with at least* 60 cells *sorted by mean for VEGFA-FLT1* ***A****. sender scores and* ***B****. receiver scores.*

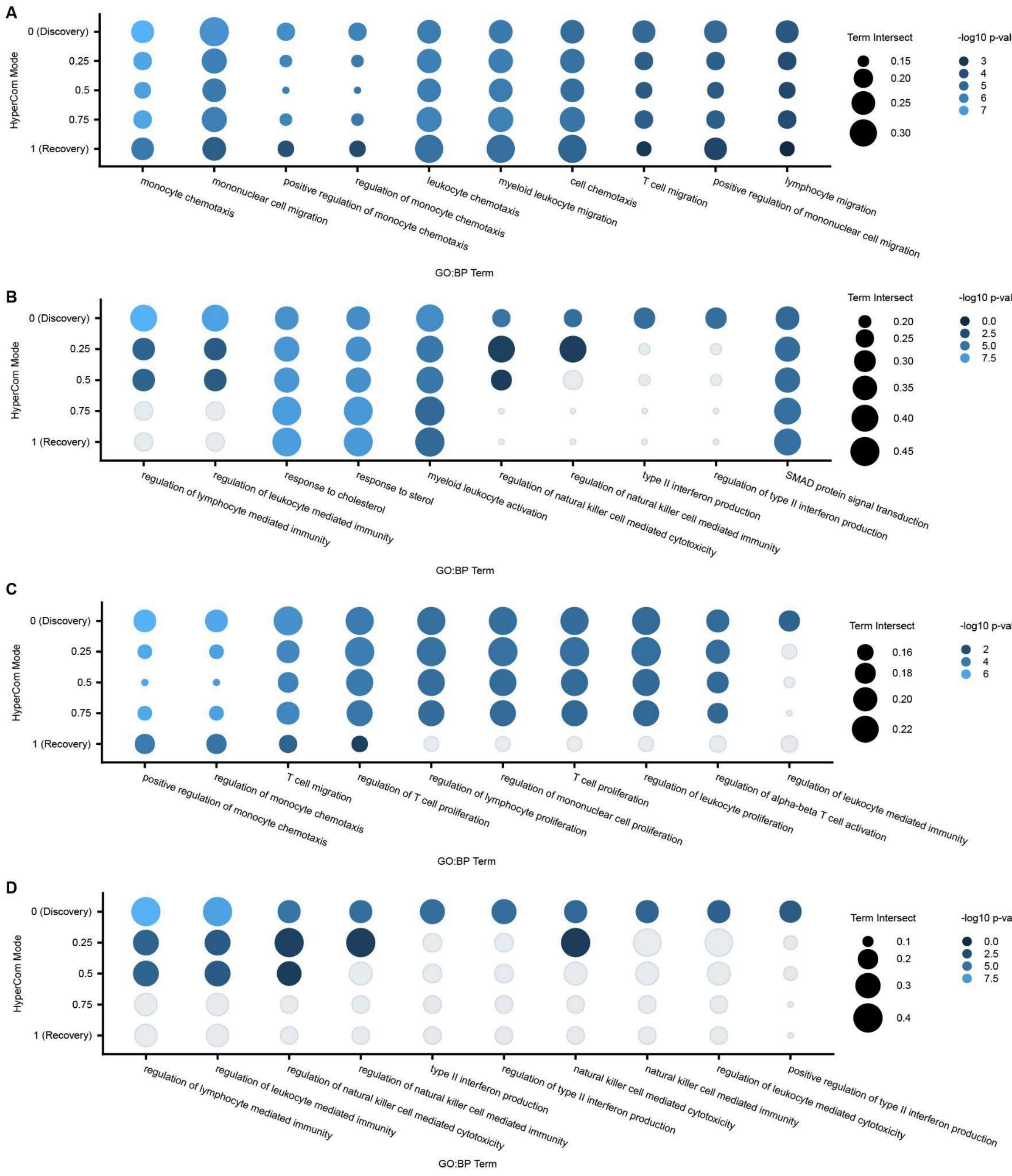


*Supplemental Figure 6.* ***GO Term Enrichment for Day 7 murine model of heart failure****. Top 10 GO:BP terms by significance for LR pair genes for a range of settings between recovery mode and discovery mode scaled by the size of the intersection with the GO:BP term and colored by -log10 p-value for genes from LR pairs with absolute mean HyperCom scores* ***A****. increasing from Sham to TAC and decreasing from TAC to BRD4KO (top 10 discovery mode terms shown, ordered by discovery mode significance from left to right)* ***B****. increasing from Sham to TAC but not decreasing from TAC to BRD4KO (top 10 discovery mode terms shown, ordered by discovery mode significance from left to right)* ***C****. increasing from Sham to TAC and decreasing from TAC*

*to BRD4KO (top 10 discovery mode terms shown, ordered by largest decrease in significance from discovery to recovery mode from left to right)* ***D****. increasing from Sham to TAC but not decreasing from TAC to BRD4KO (top 10 discovery mode terms shown, ordered by largest decrease in significance from discovery to recovery mode from left to right).*
